# Contiguously-hydrophobic sequences are functionally significant throughout the human exome

**DOI:** 10.1101/2021.09.02.458776

**Authors:** Ruchi Lohia, Matthew E.B. Hansen, Grace Brannigan

**Author notes:** **For correspondence:** (GB). These authors contributed equally to this work (Matthew E.B. Hansen, Grace Brannigan). Stanley Institute for Cognitive Genomics, Cold Spring Harbor Laboratory, USA.

## Abstract

Hydrophobic interactions have long been established as essential to stabilizing structured proteins as well as drivers of aggregation, but the impact of hydrophobicity on the functional significance of sequence variants has rarely been considered in a genome-wide context. Here we test the role of hydrophobicity on functional impact using a set of 70,000 disease and non-disease associated single nucleotide polymorphisms (SNPs), using enrichment of disease-association as an indicator of functionality. We find that functional impact is uncorrelated with hydrophobicity of the SNP itself, and only weakly correlated with the average local hydrophobicity, but is strongly correlated with both the size and minimum hydrophobicity of the contiguous hydrophobic domain that contains the SNP. Disease-association is found to vary by more than 6-fold as a function of contiguous hydrophobicity parameters, suggesting utility as a prior for identifying causal variation. We further find signatures of differential selective constraint on domain hydrophobicity, and that SNPs splitting a long hydrophobic region or joining two short regions of contiguous hydrophobicity are particularly likely to be disease-associated. Trends are preserved for both aggregating and non-aggregating proteins, indicating that the role of contiguous hydrophobicity extends well beyond aggregation risk.

**Statement of Significance:** Proteins rely on the hydrophobic effect to maintain structure and interactions with the environment. Surprisingly, no signs that amino acid hydrophobicity influences natural selection have been detected using modern genetic data. This may be because analyses that treat each amino acid separately do not reveal significant results, which we confirm here. However, because the hydrophobic effect becomes more powerful as more hydrophobic molecules are introduced, we tested whether unbroken stretches of hydrophobic amino acids are under selection. Using genetic variant data from across the human genome, we found evidence that selection pressure increases continually with the length of the unbroken hydrophobic sequence. These results could lead to improvements in a wide range of genomic tools as well as insights into disease and protein evolutionary history.

## Introduction

An ambitious task for human genetics is discovering the genetic basis for heritable traits and disease risks. In principle, genome-wide association studies (GWAS) can identify variants associated with an effect, independent of the genomic or protein context of the variant (***Visscher et al., 2017***). In practice, associated variants are often not directly causal but instead passively “tag” haplotypes containing the true causal variants that are either unknown or observed but not directly tested for association (***Cannon and Mohlke, 2018***). The pattern of linkage disequilibrium that defines “tagging” is population dependent. Any information on the protein-level function of the local sequence surrounding an associated variant can help fine-map and rank putative causal variants from a list of associations(***Gallagher and Chen-Plotkin, 2018***).

All protein level methods for variant-to-function prediction rely on some form of residue characterization. In addition to physicochemical properties (***Stone, 2005***; ***Niroula et al., 2015***; ***López-Ferrando et al., 2017***; ***Hecht et al., 2015***; ***Popov et al., 2019***) these may include evolutionary conservation (***Ng and Henikoff, 2001***; ***Thomas and Kejariwal, 2004***; ***Stone, 2005***; ***Capriotti et al., 2006***; ***Choi et al., 2012***; ***Hecht et al., 2015***; ***Niroula et al., 2015***) and structural propensities (***Capriotti et al., 2004, 2005***; ***Parthiban et al., 2006***; ***Wainreb et al., 2010***; ***Popov et al., 2019***; ***Ittisoponpisan et al., 2019***; ***Iqbal et al., 2020***). Such methods may also rely on known protein structures in order to incorporate properties such as local secondary structure and solvent accessibility (***Iqbal et al., 2020***). In the absence of structural information, however, computational prediction accuracy is low, and full three-dimensional structures have been experimentally solved for a tiny fraction of known proteins (***Rose et al., 2016***; ***Mir et al., 2017***). Fewer than 35% of human protein coding genes have structures deposited in the protein data bank (***Prlić et al., 2016***). With few exceptions, therefore, these methods have not been applied on a genome-wide scale.

Most functional proteins are intrinsically modular. For example, a given protein may include multiple secondary structure elements, ordered and disordered regions, transmembrane and soluble portions, or stretches of highly charged regions followed by stretches of highly hydrophobic regions. In the absence of structural information, bioinformatics “variant-to-function” prediction methods that use sequence context typically neglect modularity, by defining the local sequence using a window of constant length, centered around the mutation (***Teng et al., 2010***; ***Capriotti et al., 2006***; ***Capriotti and Fariselli, 2017***; ***Hecht et al., 2015***). This definition of sequence context can weaken predictive power, particularly for SNPs near the boundary between adjacent protein modules with distinct functional roles.

Despite the common framing that “sequence determines structure which determines function”, single nucleotide polymorphisms (SNPs) can alter function while leaving the protein structure essentially unchanged. For example, intrinsically disordered proteins (IDPs) lack unique structure yet are both essential for many critical biological pathways (***Ward et al., 2004***; ***Dyson and Wright, 2005***; ***Uversky, 2013, 2019***; ***Panchenko and Babu, 2015***) and sensitive to sequence. (***Weinreb et al., 1996***; ***Uversky et al., 2012***; ***Cuanalo-Contreras et al., 2013***; ***Patel et al., 2015***; ***Uversky, 2015***; ***Tovo-Rodrigues et al., 2016***).

Although not all functional proteins have well-defined structures, we showed in a previous study(***Lohia et al., 2019***) that a long intrinsically disordered protein can retain modularity of interactions. Fully-atomistic molecular dynamics simulations of a 92-residue disordered protein, run using enhanced sampling, revealed a soft network of tertiary contacts between contiguous stretches of hydrophobic residues. In order to distinguish these stretches from any other more traditional segment or domain definition, we follow terminology more common to polymer physics and call these stretches “blobs”. Blobs may contain secondary structure elements, but are not required to do so.

Here we demonstrate that hydrophobic blobs can be used as generic predictors of functional modules, by testing for enrichment of functional effects throughout the proteome. Our hypothesis was motivated by the cooperativity of the hydrophobic effect (***Jiang et al., 2017***) and the recognition that hydrophobic residues are likely to be buried (***Lins et al., 2003***) and contain a high density of interactions. Any curve drawn through such a cluster, including a curve representing the peptide chain, will contain contiguous hydrophobic residues (Figure 1).

**Figure 1.**
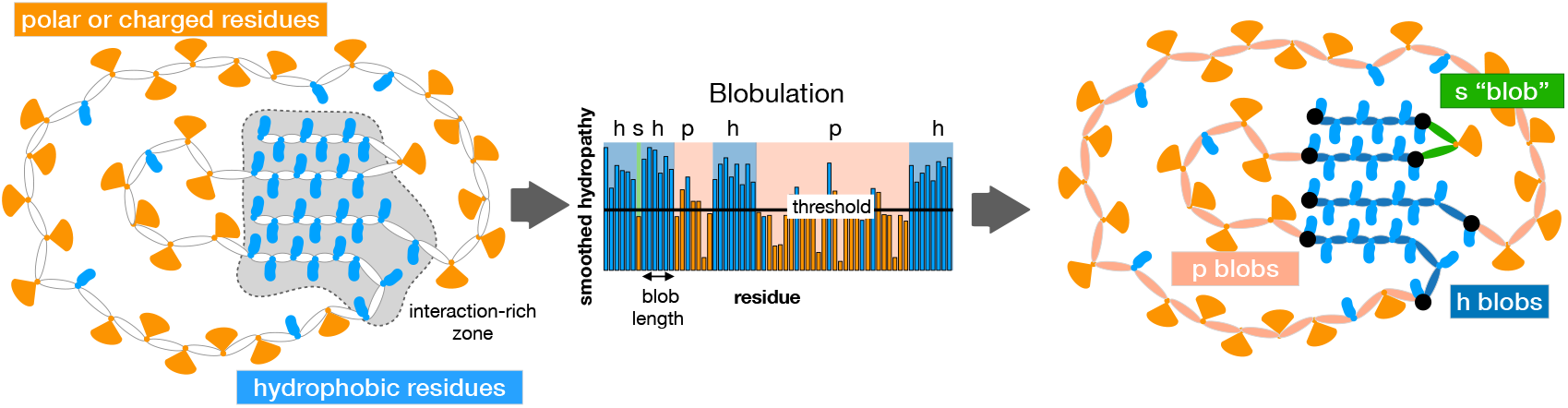
Cartoon representation of the blobulation approach. Left: A given protein chain consists of amino acids that are either hydrophobic (blue ovals) or not hydrophobic (orange fans). Clusters of hydrophobic residues (gray shading) are likely to be buried and have a high number of interacting residues. Any curve drawn through such a cluster, including the curve representing the peptide chain, will contain contiguous hydrophobic residues. Center: The blobulation algorithm calculates the smoothed hydrophobicity for each residue, and then identifies stretches of at least *L*_min_ residues above the hydrophobicity threshold (black line), where *L*_min_ = 4 unless otherwise noted. This process defines the boundaries between blobs, and also provides information about the blob hydrophobicity class: blobs of contiguous hydrophobic residues are “h” blobs (at least *L*_min_ residues), while remaining stretches are either p “blobs” (at least *L*_min_ residues) and “s blobs” (fewer than *L*_min_ residues). Right: Blobs mapped onto the peptide chain; the peptide backbone is colored according to blob hydrophobicity class and black circles mark the boundary between blobs. In this illustration there are 4 h-blobs, 2 p-blobs, and 1 s-blob.

In the present study, we used a large database of protein sequences to determine the extent to which hydrophobicity-based sequence segmentation captures functional modules. Our test involves three steps: 1) segmentation of the protein into the blobs (“blobulation”), 2) characterization of the blobs based on various physicochemical properties and 3) testing whether SNPs with “known” functional impact are enriched or depleted genome-wide in blobs with particular properties. We test for enrichment of hydrophobic blobs among disease-associated SNPs, repeating the calculation with a range of hydrophobicity thresholds and stratifying according to hydrophobic blob length. This process reveals a striking dependence that is muted when a standard moving window approach is used instead, and is completely lost when the sequence context is neglected. We test for consistency against population frequency data, finding that disease-associated SNPs in extremely hydrophobic blobs are under greater purifying selection than those in less hydrophobic blobs. We further test the functional effects of mutations that split or merge blobs or change other categorical properties. In doing so, we find that blobulation yields a meaningful protein topology that is useful for sequence characterization beyond hydrophobicity.

## Results

### Regions of contiguous hydrophobicity are enriched for disease-causing SNPs

In order to test whether the hydrophobicity of a reference allele was correlated with its functional impact, we calculated the enrichment of dSNPs relative to nSNPs as a function of hydrophobicity of the reference allele. As shown in Figure 2b, we did not detect any significant correlation (Pearson’s r=0.02, n=17). Hydrophobicity is a highly cooperative phenomenon (***Jiang et al., 2017***), and we hypothesized that the hydrophobicity of the local sequence could be critical for determining the functional effects of the dSNP. We considered two methods for determining local sequence context: our own “blobulation” approach and a fixed-width moving window.

**Figure 2.**
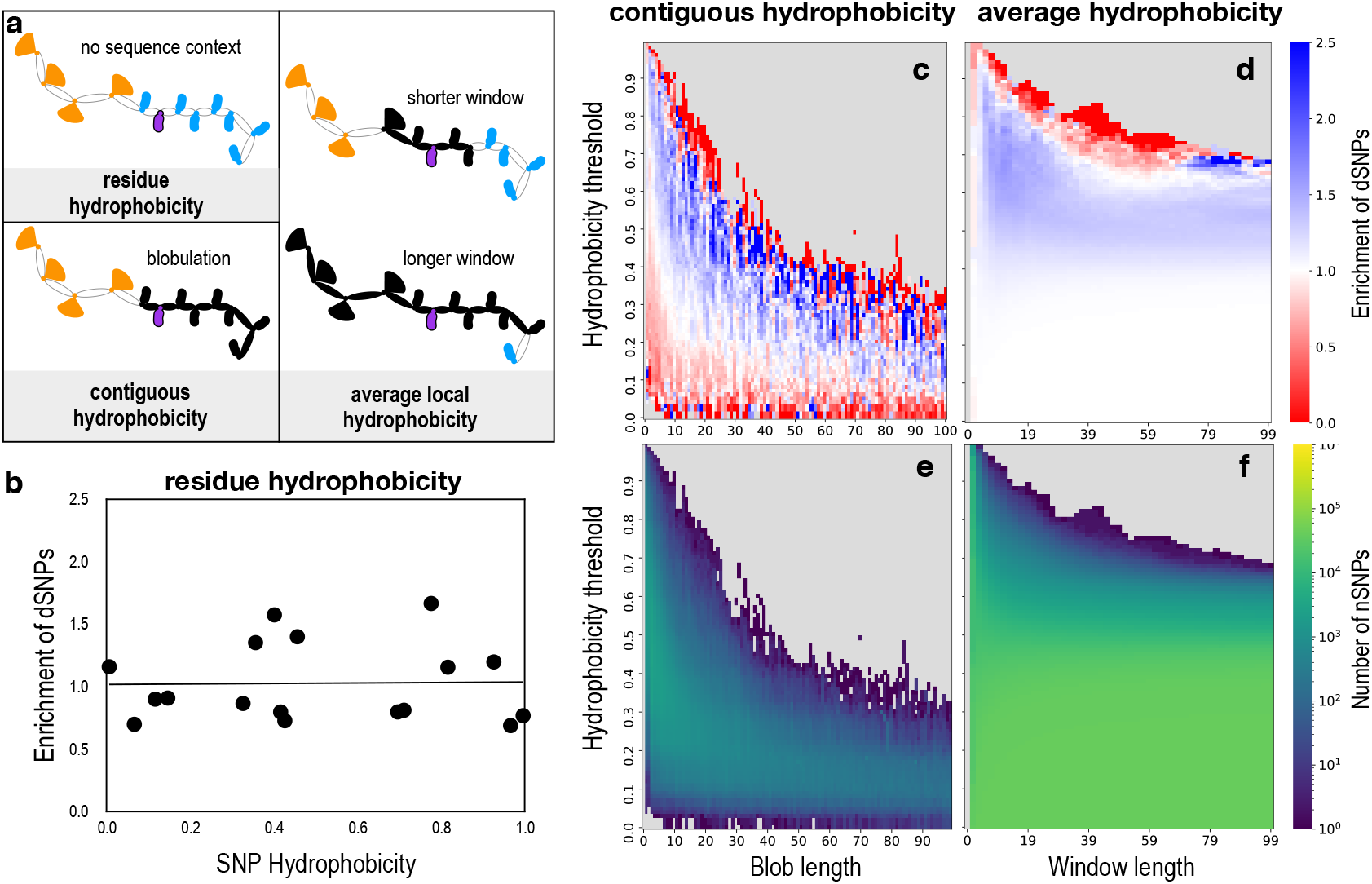
Effect of segmentation approach, length, and hydrophobicity threshold on calculated enrichment of dSNPs in hydrophobic segments. a) Illustration of three measures of SNP hydrophobicity (residue, contiguous, and average) for the amino acid shown in purple, found within a hypothetical peptide chain where hydrophobic residues are blue ovals and non-hydrophobic residues are orange fans. The local sequence determined by each segmentation method is shown in black fill. b) Enrichment of dSNPs relative to nSNPs as a function of hydrophobicity of the reference allele, with line of best fit. No trend or significant correlation is observed (Pearson’s r=0.02, p=0.94, n=17). c-d) Enrichment of dSNPs relative to nSNPs for hydrophobic segments for varying hydrophobicity and length thresholds using a blobulation approach (c) and moving window approach (d). Bins with no data are shown in gray. e-f) The number of nSNPs in hydrophobic segments per bin using the blobulation approach (e) and moving window approach (f).

Blobulation (Figure 1) is a particular systematic approach for identifying modular components in the protein sequence, which we previously introduced (***Lohia et al., 2019***) for analysis of a specific protein. Here we have generalized and extended the approach by varying two previously-fixed parameters (minimum blob length and hydrophobicity threshold) that are required to define blob edges, as described in *Methods*. The degree of enrichment for disease-association is expected to be dependent upon both of these parameters, and we ran enrichment tests for a wide range of minimum blob lengths and hydrophobicity threshold values. Unless otherwise noted, dSNPs are tested for enrichment relative to the expectation set by nSNPs. For example, the phrase “dSNPs are enriched in X blobs” means that dSNPs are found at a higher rate in blobs of type X than are nSNPs.

We found a consistent trend relating hydrophobicity of the local blob and enrichment of disease-associated SNPs (Figure 2 C). dSNPs are depleted in weakly hydrophobic blobs, neutral for moderately hydrophobic blobs, and become more enriched as the blob gets longer and/or satisfies a stricter hydrophobicity threshold. The trend is steadily monotonic, which supports the hypothesis that hydrophobic blobs constitute identifiable functional elements. The exception is at the plot boundary: surprisingly, blobs that satisfied the very strictest criteria were depleted in dSNPs. In the next section, we consider two potential reasons for this depletion: either dSNPs in these blobs are so deleterious that they are selected out of the population, or nSNPs in these blobs are functional as expected, but not deleterious. In addition to a consistent trend, the analysis returns a large spread in enrichment/depletion values: 3.2% of the bins in Figure 2c have a significant enrichment of less than 0.5, while 13% have a significant enrichment of greater than 1.5, and 3.5% have a significant enrichment greater than 3 (Data set 1). This result indicates that hydrophobicity-based sequence segmentation could be particularly useful for assessing the riskiness of SNPs located in long and very hydrophobic sequences.

In order to compare the effectiveness of blobulation to a moving window approach, we ran analogous enrichment tests for sliding windows across all protein sequences. Two analogous parameters (window length and mean hydrophobicity threshold) are used for defining hydrophobic regions in the moving window approach, with the resulting enrichment for a range of these values also shown in Fig 2d. Compared to the blobulation approach, the enrichment detected using the moving window approach is much less sensitive to segment length for windows beyond 6 residues (consider the tilted color bands for the blobulation approach vs the horizontal color bands for moving windows in Figure 2c-d). The significant depletion in dSNPs at the most hydrophobic edge is also preserved in the moving window analysis.

While there is no “standard” window size, most SNP prediction programs use a window size in the range of 1-21 residues (***Teng et al., 2010***; ***Capriotti et al., 2006***; ***Capriotti and Fariselli, 2017***; ***Hecht et al., 2015***). The window size is chosen to balance concerns that small window sizes may not accurately capture the “local” sequence (***Chen et al., 2006***; ***Schlessinger and Rost, 2005***; ***Sander et al., 2006***) whereas long window sizes can increase signal to noise ratio (***Park et al., 2007***). In either case, the use of a fixed-width window neglects the inherent dispersion in the size of protein modules (Figure 2 a). The enrichment insensitivity to window size was thus surprising, but the expected noise was likely lost due to averaging across the proteome. As shown in Fig 2e, the overall fraction of hydrophobic segments for mild or moderate hydrophobicity thresholds was also insensitive to window size, confirming that the data set we use here is sufficiently large to average out large window noise. For a single SNP, considering too many residues far from the SNP will still contribute significant noise to the prediction.

Overall, use of the moving window approach yields two-fold enrichment for only a few, scattered parameter values with very few qualifying sequences. In contrast, blobulation yielded greater than two-fold enrichment for a well-defined set of parameters (hydrophobicity threshold > 0.4 and minimum blob size > 15). This observation supports our hypothesis that blobulation provides a more meaningful and less noisy approach to protein segmentation than use of a fixed-length moving window.

### Population frequency signatures of differential selection on hydrophobic blobs

If hydrophobic blobs capture functional segments of coding regions, then we may expect to see signatures of differential selective constraint with varying blob properties. We probe this by examining the genetic diversity of SNPs, stratified by their surrounding blob hydrophobicity. A benchmark measure of genetic diversity is the expected heterozygosity Θ ≡ 2*v*(1 − *v*), where *v* is the frequency of the coded allele. Sequences under more functional constraint, like exons in essential proteins, experience purifying selection (removal of almost all new functional alleles), which lowers Θ compared to regions under little or no constraint (see, e.g. ***Dickerson (1971***); ***Charlesworth et al. (1993***); ***Sunyaev et al. (2000***); ***Siepel et al. (2005***); ***Somel et al. (2013***)). Conversely, balancing selection (maintenance of multiple functional alleles) causes increased genetic diversity over a genomic region (see, e.g. ***Llaurens et al. (2017***)). Population substructure can also increase the genetic diversity, but such effects would occur genome-wide and would not be correlated with blob hydrophobicity class.

For population frequency data we use gnomAD v3 SNP frequency data for *N* ∼ 32, 000 non-Finnish Europeans ***Karczewski et al. (2021***). The average heterozygosity of non disease-associated SNPs over this cohort is ⟨Θ⟩ _nSNP_= 0.14, while that of disease-associated SNPs is nearly 50 times lower, ⟨Θ⟩_dSNP_ = 0 0029.. Figure 3 shows the mean heterozygosity of nSNPs and dSNPs binned by the maximal blob hydrophobicity (*H*_max_) for each SNP (see Methods). The null background is generated by random permutation of the heterozygosity assigned to each SNP, corresponding to the null hypothesis that there is no correlation between SNP heterozygosity and blob properties. Both nSNPs and dSNPs show a significant departure (outside the 1^st^ − 99^th^ null percentiles) in mean heterozygosity from null expectations for the largest *H*_max_ bin, *H*_max_ ≥ 0.75. For nSNPs, the heterozygosity is larger than expected in the highest *H*_max_ bin (> 99^th^ null percentile), consistent with relaxed purifying selection or increased balancing selection. In contrast, for dSNPs the heterozygosity in the largest *H*_max_ bin is smaller than expected under the null (< 1^st^ null percentile). Given that disease risk alleles are almost always “new” deleterious mutations, we surmise that this decrease in Θ is due to greater purifying selection acting to remove the risk allele. In other words, the disease-associated alleles in highly hydrophobic blobs appear to be more deleterious than disease-associated alleles in lower hydrophobicity blobs.

**Figure 3.**
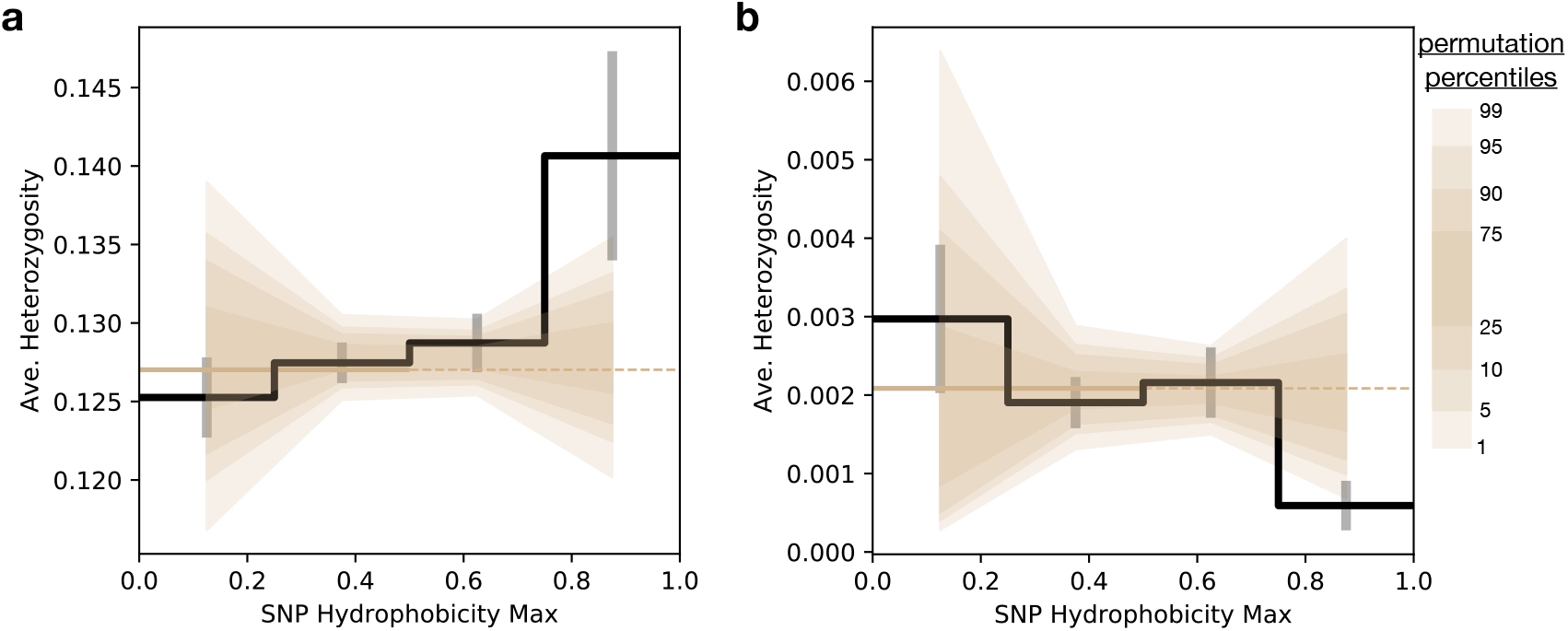
Expected heterozygosity of SNPs in Europeans as a function of blob hydrophobicity. The average expected heterozygosity Θ for SNPs in h-blobs as a function of the maximum SNP hydrophobicity *H*_max_ (binned in bin widths of Δ*H*_max_ = 0.25) is shown by the black line, with standard errors in the mean indicated. Only SNPs where at least one of two alleles is in an annotated h-blob are considered. Frequencies are taken from the gnomAD cohort of non-Finnish Europeans. The brown line is the average heterozygosity for all SNPs with *H*_max_ < 0.5 (the line is solid for *H*_max_<0.5 and dashed for *H*_max_ > 0.5). The background null distribution generated from randomized permutations are shown in brown, with the null percentiles as shown in the legend on the right. Panels a) and b) show the results for nSNPs and dSNPs, respectively.

The results here may explain the non-monotonic behavior of the dSNP-to-nSNP enrichment seen in Figure 2c and d for both the blobulation and moving windows calculations. The thin band of “red” over the bins with the highest hydrophobic threshold that still contain data indicates a depletion in the number of dSNPs relative to nSNPs. Before examining the population frequency data, it was not clear if the lack of dSNPs relative to nSNPs is due to increased selection against dSNPs in those blobs, increased balancing selection for nSNPs in those blobs, or a mixture of both. In this section we have shown that the heterozygosity of dSNPs decreases with higher blob hydrophobicity while the heterozygosity of nSNPs rises with blob hydrophobicity. This suggests that the depletion observed in Figure 2c and d in the high hydrophobicity regions is caused by both increased selection against disease-associated alleles and increased selection for the maintenance of functional non disease-associated alleles.

We tested whether the trends observed in Figure 3 are specific to non-Finnish Europeans by performing the same analysis on the gnomAD East Asian frequency data set of approximately ∼ 1, 500 individuals. Although we do not observe the same level of statistical significance, we do find the same trends as in the European cohort: for nSNPs there is increasing heterozygosity with increasing *H*_max_ and for dSNPs the heterozygosity decreases with increasing *H*_max_ (Figure S1). The stratification of genetic diversity with blob properties appears to be a feature shared across diverse human populations.

In order to provide some context for the proteins containing the blobs in the *H*_max_ ≥ 0.75 bin we use gene-ontology and pathway enrichment tests. The nSNPs found in blobs with *H*_max_ ≥ 0.75 are distributed across a subset of proteins (*n* = 635), which we test against the background of all proteins containing nSNPs (*N* = 10, 406). For the dSNPs, there are *n* = 179 proteins containing the *H*_max_ ≥ 0.75 SNPs and a total background of *N* = 1803 proteins containing a dSNP. Both the *H*_max_ ≥ 0.75 nSNP and dSNP proteins are most enriched for being located in or on a membrane (Data sets 2 and 3), as expected for proteins containing highly hydrophobic blobs. In terms of molecular function and biological process, however, the nSNPs and dSNPs differ in their enriched ontologies. The *H*_max_ ≥ 0.75 nSNPs are enriched for olfactory processes, chemical sensing, and signal transduction. This accords with previous observations that olfactory-related genes exhibit signs of relaxed purifying selection relative to other genic regions ***Pierron et al. (2013***). This analysis has essentially re-identified this same feature of greater diversity in chemical sensing pathways, found here because the increased genetic diversity is specifically present in highly hydrophobic subdomains of those proteins. In contrast, the *H*_max_ ≥ 0.75 dSNPs are enriched for cation and small molecule trans-membrane transport proteins. These SNPs reside in proteins enriched for critical membrane bound transporters, consistent with the signal of higher functional constraint.

### dSNP enrichment emerges in shorter and more weakly-hydrophobic blobs for aggregating proteins

Side-chain hydrophobicity plays a well-established role in diseases involving aggregating proteins. (***Conchillo-Solé et al., 2007***; ***Fang et al., 2013***; ***Tartaglia and Vendruscolo, 2008***). In order to test whether the trends observed in Fig 2b are amplified in aggregating proteins, we separated aggregating proteins from the primary data set. Proteins involved in the formation of extracellular amyloid deposits or intracellular inclusions with amyloid-like characteristics are included in the “Aggregating proteins” (28 proteins, 124 nSNPs, 330 dSNPs) subset, while all remaining proteins are labelled as “Non-aggregating proteins”.

Figure 4a shows the blob-type distribution of nSNPs and dSNPs, for both aggregating and non-aggregating proteins, using a mild hydrophobicity cutoff of 0.4. For dSNPs, the blob distributions are insensitive to aggregating-tendencies of the protein: a given dSNP in an aggregating protein is as likely to be found in an h-blob as if it were in a non-aggregating protein. We do still detect a small difference in enrichment because nSNPs in aggregating proteins are found slightly less frequently in h-blobs *vs* nSNPs in non-aggregating proteins.

**Figure 4.**
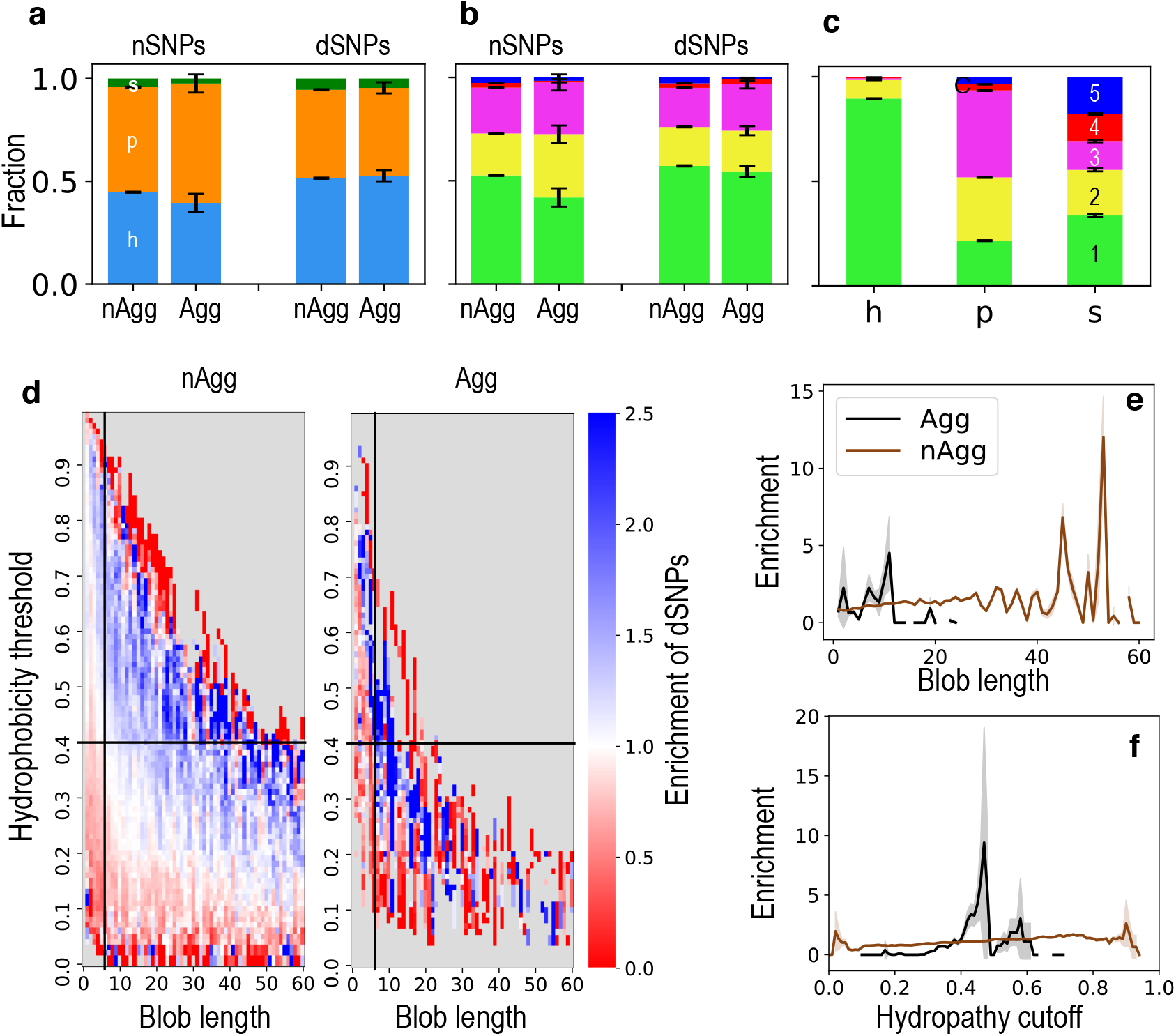
Blob hydrophobicity for SNPs in aggregating proteins. a) Fraction of nSNPs or dSNPs that are found in either blob type (a) or charge class(b) in non-aggregating proteins and known-aggregating proteins. c) is the fraction of charge classin each blob type in all proteins. The distribution in a, b and c are plotted with a cutoff and minimum blob length of 0.4 and 4 respectively. The effect of these parameters on proportion of SNPs in blob type is further discussed in Fig 2b. Error bars represent one standard error for multinomial distributed data. d) is same as Fig 2b but for non aggregating proteins and aggregating proteins. Enrichment of dSNPs in h-blob at fixed cutoff of 0.4 (e) or fixed blob length of 6.

Stratifying the enrichment by hydrophobicity cutoff and blob length (as in Fig 2b of the main text) reveals a more informative quantitative shift. Figure 4d compares the data from Fig 2b) with the same analysis for aggregating proteins. The highly-enriched (blue) band is shifted toward the origin for aggregating proteins, indicating that sensitivity to mutation is found in shorter and more weakly hydrophobic blobs. This shift is also visible in transects of the data along one row and one column, shown in Figure 4e and f. These results are consistent with previous observations that the in vitro aggregation rate of unstructured polypeptide chains is proportional to hydrophobicity and inversely proportional to net charge(***DuBay et al., 2004***).

To test the significance of charge, blobs were also assigned charge class (Figure 4b), which is the predicted globular phase according to Das and Pappu(***Das and Pappu, 2013***) and calculated using the fraction of positive and negative charges. Possible values of the blob charge class are 1 (Weak polyampholyte), 2 (Janus or boundary region), 3 (Strong polyampholyte), 4 (Negatively-charged strong polyelectrolyte), and 5 (Positively-charged strong polyelectrolyte). Sequences in class 1 have a low fraction of positively charged residues, and nearly all structured proteins fall in class 1. In contrast to structured proteins, IDPs can be found in all five Das-Pappu phases, particularly classes 3, 4, and 5. The blob hydrophobicity class and charge class are fundamentally correlated; while blob charge class does not explicitly consider hydrophobicity, increasing the number of charged residues will reduce the average hydrophobicity of a blob. The extent of this correlation is shown in Figure 4C, which breaks down the fraction of h and p blobs that fall in each Das-Pappu charge class. As expected, most h blobs (90%) fall in class 1 (weak polyampholyte), followed by 9% in class 2 (Janus). The p blobs are more evenly distributed across classes, with the highest fraction (42%) classified as strong polyelectrolytes.

Since there are five charge classes and only three hydrophobicity classes, we hypothesized that blob charge class would be more precise than blob hydrophobicity class for identification of functional protein segments. Instead, we found that the maximum charge class enrichment (1.09 fold for weak polyampholyte blobs) is comparable or even slightly less than the enrichment found for hydrophobicity class (1.15 fold). This result suggests that even short contiguous stretches of weak hydrophobicity are as effective as charge distribution for predicting functional impact. In the context of aggregating proteins, classification by charge yielded similar patterns as classification by hydrophobicity: dSNPs have equivalent distributions regardless of protein aggregation tendency, while nSNPs in aggregating proteins are significantly less likely to be found in globular, weak-polyampholyte blobs (Class 1) than those in non-aggregating proteins.

### Disease-associated SNPs are enriched for mutations that change local blob characteristics and overall protein blob topology

The blobulation process yields a series of h blobs, connected by p and s linkers, which we term the “blobular topology”. Such a topology is analogous to the classic protein topology of secondary structure elements. A SNP can alter the blobular topology by moving a short stretch of contiguous residues above or below the minimum blob size, either forming a new small h-blob or dissolving an existing small h-blob respectively. In the background case we expect to see more formation than dissolution, since the blob frequency decreases with length (Figure 2 e), and there are more blobs just below the minimum length than just above it. Figure 5A confirms this expectation for nSNPs. A SNP can also split a long h-blob by interrupting a long contiguous hydrophobic sequence, or merge two smaller h-blobs into one long h-blob by removing such an interruption. Using the mild hydrophobicity threshold of 0.4 and minimum blob length of 4 residues, fewer than 15% of SNPs overall cause a topological change. As shown in Figure 5C, however, certain topological changes are strongly enriched in dSNPs. While the difference between the fraction of nSNPs and dSNPs that form h-blobs is not significant (*p* > 10^−3^), dSNPs are significantly more likely to dissolve h-blobs (*p* < 10^−21^), split h-blobs (*p* < 10^−80^), and merge h-blobs (*p* < 10^−36^). The magnitude of enrichment (2.1-fold) is greatest for SNPs that split longer h-blobs into two shorter ones and is only moderately weaker for the reverse merging of two h-blobs (1.7-fold). These results are consistent with the functional sensitivity of long blobs shown in Figure 2c.

**Figure 5.**
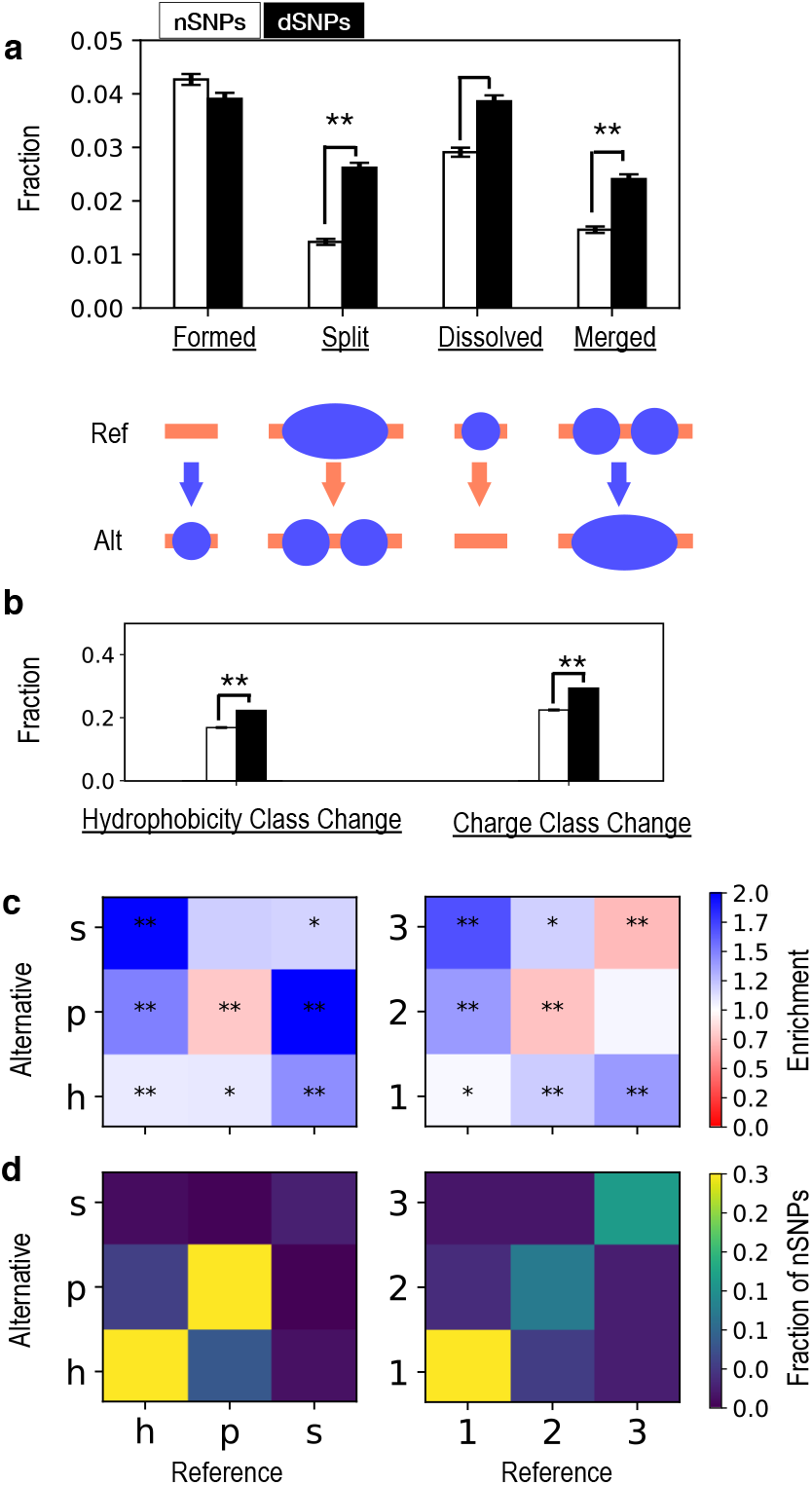
Distribution of SNPs that change blob properties. a) Fraction of nSNPs or dSNPs that change the blobular topology by either forming or dissolving an h-blob or by splitting one h-blob or merging two h-blobs. b) Fractions of nSNPs or dSNPs that induce a change in hydrophobicity class or charge class. c) The enrichment of dSNPs relative to nSNPs that induce a specific blobular topology transition between the reference allele (x-axis) and the alternative allele (y-axis). This is shown for two blob properties, hydrophobicity class (left) and charge class (right). d) The overall proportion of nSNPs that induce each of the blobular topology transitions shown in c). Fewer than 1.5% of SNPs involve charge classes 4 or 5, and these data are not shown for readability. Significant enrichment or depletion in dSNPs is annotated with one star (p < 10^−3^) or two stars (p < 10^−10^). Errors bars in (a) and (b) represent one standard error for multinomial (a) or binomial (b) distributed data. In all cases blobulation is calculated using *H*^⋆^ = 0.4 and *L*_min_ = 4.

Regardless of overall topological changes, SNPs may change the characteristics of their local blob. Such changes may affect blob topology (as in Figure 5a), or simply shift blob boundaries (as in Figure 5b). We now broaden the analysis to include changes in blob characteristics that do not necessarily affect the blob boundaries. We observe that about 17% of the nSNPs and 22% of dSNPs introduce a blob-hydrophobicity class change, yielding a 1.3 fold enrichment for blob hydrophobicity class changes (Fig 5b). Fig 5c compares the frequency of possible blob hydrophobicity class transitions. Mutations involving h → p blob transitions yield the maximum enrichment (1.5 fold) among dSNPs. In contrast, dSNPs are only 1.1 fold enriched in the reverse transition (p → h), and SNPs that remain in p blobs for both the ancestral and derived allele in p blobs are depleted among disease-associated SNPs.

Similarly, the frequency of SNP-induced changes in blob charge class is shown in Fig 5c, for transitions involving blobs in categories 1, 2, and 3. Transitions involving categories 4 or 5 (positively or negatively-charged strong poly electrolyte) represented fewer than 1.5 % of the total transitions. Disease-associated SNPs are enriched for all mutations that change blob charge class and either unenriched or weakly depleted for mutations that do not change the local blob charge class. so 1 → 3 > 1 → 2 and 3 → 1 > 3 → 2. Collectively, these results are consistent with the increased mutational sensitivity of hydrophobic (and typically buried) blobs that is shown in Figure 2, while also emphasizing that mutations that *change* charge class are particularly likely to be causal. For instance, charge reversal of a charged residue could have particularly strong functional effects. While these effects might be amplified if the residue was located in an h-blob (and thus interacting with other protein residues), the charge reversal itself would not affect the blob hydrophobicity class.

## Discussion

In the present work, we have presented the “blobulation” scheme for identifying interaction-rich protein regions from sequence, and tested its use in detecting functional modules across the proteome. We find that enrichment of disease-associated mutations in hydrophobic blobs increases with the strictness of the hydrophobic blob criteria, with greater than 4-fold enrichment for disease-association in the longest, most hydrophobic blobs. The range and resolution of varying enrichment is strongly damped in the *status quo* fixed-length moving window approach. Stratifying SNPs by their surrounding blob properties reveals genome-wide differences in blob genetic diversity, demonstrating pervasive differential selection on at least some classes of blobs.

We also find that disease-associated mutations are significantly more likely than non-disease associated mutations to change the topology of the sequence. This suggests that blobulation provides a meaningful topology that can be used as a framework for sequence analysis, and which requires only the protein amino acid sequence and two parameters (minimum blob length and hydrophobicity threshold). Once blobs are identified, they can be characterized on any property of interest. As an example, in the present work, we also find that disease-associated mutations are moderately enriched for mutations that cause transitions in blob hydrophobicity class (up to 1.5 fold) and strongly enriched for mutations that cause certain transitions in blob charge class (1.7 fold).

While we are not aware of a similar approach applied to generic proteins, hydrophobic blobs are analogous to the aggregation “hot spots” identified by tools such as AGGRESCAN (***Conchillo-Solé et al., 2007***), ProA (***Fang et al., 2013***), and Zyggregator (***Tartaglia and Vendruscolo, 2008***). We do find that functional sensitivity in aggregating proteins follows similar trends but emerges in shorter blobs satisfying weaker hydrophobicity criteria. The difference between aggregating and generic proteins is thus quantitative, not qualitative, and demonstrates that hydrophobic interactions occur on a useful continuum.

Structural data implicitly provides insight into the segmentation of a protein sequence, but we show here that segmentation may be meaningfully estimated from the sequence alone. We have proposed blobulation as a generalizeable approach for segmentation, although in principle, secondary structure prediction could be used instead. For many proteins, however, this approach would be frequently unfeasible and overly restrictive. In addition to the challenges inherent in predicting secondary structure (which may be highly sensitive to the local protein environment or simply non-existent), many secondary structure prediction methods require alignment to a homologous sequence with known structure (***Yang et al., 2018***; ***Zhang et al., 2018***; ***Wang et al., 2017***; ***Ma et al., 2018***). This is essential for secondary structure predictors to achieve their primary goal: distinguishing *between* secondary structures. Segmentation, however, only requires knowing where the segment begins and ends.

We anticipate that these results may further efforts to assess functional significance of SNPs for which disease-causality is uncertain. Despite the wealth of genetics information presently available, the complex nature of many phenotypes makes the identification of causal variants difficult (***Claussnitzer et al., 2020***). SNP-prediction methods often test residue properties for relevance by machine learning algorithms, which derive their decision rules based on training datasets of annotated mutations, and therefore are sensitive to the training set used. Machine learning methods also rely on a large number of features, which can obscure the biological relevance of any given feature. Here we have both performed hypothesis-driven tests for enrichment of certain features in putatively causal SNPs and provided a two-dimensional table (Dataset 1) of wide-ranging enrichments as a function of blob properties, which can be used as inputs in future predictions.

## Methods and Materials

### SNP data

The SNP data we use is from the UniProtKB literature curated list of missense variants (www.uniprot.org/docs/humsavar, obtained on 17-Jun-2020) (***Yip et al., 2008***). Variants are annotated using the American College of Medical Genetics and Genomics/Association for Molecular Pathology (ACMG/AMP) terminology(***Richards et al., 2015***). dSNPs are those annotated as “likely pathogenic or pathogenic” (*N* = 30, 227), and nSNPs are those annotated as “likely benign or benign” (*N* = 39, 448).

### Blobulation

Blobulation refers to the partitioning of a given amino acid sequence (e.g. a protein isoform) into three kinds of segments based on the local hydrophobicity. The segments are referred to as “blobs”, and they are assigned a primary type known as the “hydrophobicity class”. The three blob hydrophobicity classes are hydrophobic (“h-blob”), non-hydrophobic or polar (“p-blob”), and short non-hydrophobic spacers (“s-blobs”). We use a simple partitioning approach, described below, that is based on three parameters: the h-blob hydrophobicity threshold *H*^⋆^, the minimum length for hydrophobic segments to be labeled as an h-blob 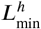, and the threshold length separating p-blobs from s-blobs,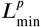.

Every amino acid *i* is assigned a mean hydrophobicity *H*_*i*_, defined as the average Kyte-Dolittle (***Kyte and Doolittle, 1982***) score with a window size of three residues, scaled to fit between 0 and 1. Given the hydrophobicity threshold *H*^⋆^, we first identify all contiguous stretches where *H*_*i*_ ≥ *H*^⋆^ for all *i* in the sub-sequence and where the sub-sequence length *x* is 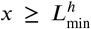; these are classified as h-blobs. The sub-sequence between a pair of h-blobs, or between an h-blob and the end of the amino acid sequence (for sub-sequences at an edge), is classified as a p-blob if the length 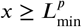, and is classified as an s-blob if 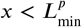. In all analyses used in this work, we set 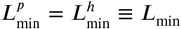 so that there are effectively only two blobulation parameters, *H*^⋆^ and *L*_min_.

Consider a given protein amino acid sequence that contains a missense SNP at residue *i*. For a missense SNP, the reference allele and the alternative allele correspond to two different amino acids. We index the two amino acids for this polymorphism by *α* ∈ {R, A}, where R is the amino acid corresponding to the reference allele and A for the alternative allele. Upon blobulation of this sequence with amino acid *α* at *i*, the residue *i* will be contained within a blob type *β* ∈ {*h, p, s*} with some length *x*. We define the binary blobulation query function *B*_*α,β,x*_(*i*)

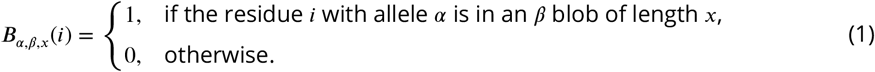

Importantly, *B*_*α,β,x*_(*i*) is dependent upon the hydrophobicity threshold *H*^⋆^ (as are all quantities that depend on *B*_*α,β,x*_(*i*)). Furthermore, because h-blobs are detected first, p-blobs are detected from the remaining residues, and s-blobs are assigned last, *B*_*α,p,x*_(*i*) will be dependent upon the minimum h-blob length, and *B*_*α,s,x*_(*i*) will be dependent upon the minimum h and p-blob lengths. Unless stated otherwise, blobulation is based on the reference allele for all SNPs and we suppress the explicit *α* notation.

### Das-Pappu charge class

The blob charge class is a secondary blob property representing the Das-Pappu phase (***Das and Pappu, 2013***), which is determined using the fraction of positively and negatively charged residues. Blobulation does not rely on charge class, but any blob may be assigned a charge class following blobulation. Possible values of the blob charge class are 1 (Weak polyampholyte), 2 (Janus or boundary region), 3 (Strong polyampholyte), 4 (Negatively-charged strong polyelectrolyte), and 5 (Positively-charged strong polyelectrolyte).

### dSNP enrichment

Let *f*_*β,x*_ and *g*_*β,x*_ denote the fraction of nSNPs and dSNPs, respectively, which are found in *β*-type blobs of length *x*:

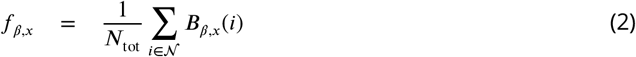

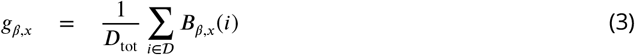

where 𝒩 is the set of nSNPs with size *N*_tot_ ≡ ‖𝒩‖ and 𝒟 is the set of dSNPs with size *D*_tot_ ≡ ‖𝒟‖. Enrichment of dSNPs relative to nSNPs for *β*-blobs of length *x* is then

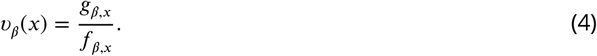

For some comparisons we consider the enrichment over all blob lengths above some minimum threshold 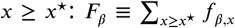 and 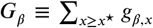 for nSNPs and dSNPs, respectively. The corresponding minimum-length enrichment is ϒ_*β*_ = *G*_*β*_ /*F*_*β*_. For the heat-map in Figure 2 we consider the enrichment over SNPs at a given hydrophobicity threshold *H*^⋆^ for lengths within a length bin. Within a length bin [*x*_0_, *x*_1_], the fractions are summed over all *x* within that bin: 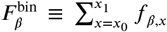 for nSNPs and similarly for dSNPs. The enrichment is given by the ratio of these bin fractions, 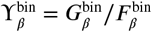.

Enrichment values are tested for statistically significant deviations from the null expectation of *ν* = 1 using a Binomial distribution for *N*_trials_ = *D*_tot_ Bernoulli trials with probability of success *p* = *f*_*β,x*_ per trial, where the expected number of “successes” is *D*_tot_ *f*_*β,x*_. The minimum-length enrichment ϒ, and enrichment within length bins, are tested similarly. Reported p-values correspond to the two-sided test, and any enrichment or depletion in dSNPs is termed “significant” for p-value < 10^−3^.

### Blob transitions induced by a SNP

For each protein sequence and each SNP *i* in the given sequence, the blobulation algorithm is performed separately for both the reference and the alternative allele. If the protein contains multiple SNPs, then the reference allele is used for all residues except *i*, i.e. we do not consider multi-SNP *cis* effects on blobulation. In order to test the statistical significance of the observed transition rates for dSNPs, we compare the observed rates to those for nSNPs. Specifically, suppose the observed fraction of nSNPs that induce a transition from blob class *β*_R_ for the reference allele to blob class *β*_A_ for the alternative allele is *f*(*β*_R_, *β*_A_), and that the fraction of dSNPs that induce such a transition is *g*(*β*_R_, *β*_A_). Then the number of Bernoulli trials is *N*_trials_ = *D*_tot_ with probability of success *p* = *f*(*β*_R_, *β*_A_), where the expected number of “successes” is *D*_tot_*f*(*β*_R_, *β*_A_) and the observed number of successes is *D*_tot_ *g*(*β*_R_, *β*_A_). Reported p-values correspond to the two-sided test, and any enrichment or depletion in dSNPs is termed “significant” for p-value < 10^−3^.

### Fixed-length moving windows

Calculations involving fixed-length moving windows apply a threshold test to the mean hydrophobicity across the window, rather than testing each residue. The mean hydrophobicity of a fixed-length moving window of length *x* (for odd *x*) centered around residue *i* is:

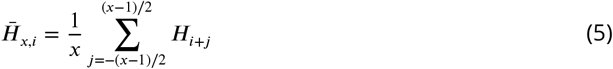

and the associated query function for the moving-window equivalent of h-blobs (*β* = *h*) with the reference allele at *i* (*α* = R) is:

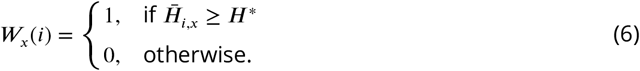

The query function *W* is used in place of *B* for Equations 2 and 3. All subsequent equations remain unchanged.

### Population frequency data

Frequency data is from the gnomAD v3 genomes dataset of variants for the non-Finnish European cohort: gnomad.genomes.r3.0.sites.chr*.vcf.bgz, accessed July 24, 2020, using the “INFO/AF_nfe_male” and “INFO/AF_nfe_female” tags. This cohort contains 32, 299 individuals, providing allele frequency data as low as ∼ 1/60, 000 ∼ 0.002%. Variants with dbSNP identifications (“rsids”) were intersected with the UniProt SNP data. There are 36,025 SNPs in common (in 10,565 genes), composed of 29,653 nSNPs (in 10,143 genes) and 6,372 dSNPs (in 1,594 genes). For comparison we also analyzed the gnomAD v3 East Asian cohort, using the “INFO/AF_eas_male” and “INFO/AF_eas_female” tags. This cohort contains 1, 567 individuals.

### Expected heterozygosity of SNPs in h-blobs

We test for the impact of the blob hydrophobicity on SNP heterozygosity by comparing the average expected heterozygosity to the null distribution in different hydrophobicity bins, described as follows. We first partition SNPs by the maximum blob hydrophobicity threshold for which at least one of the alleles remains in an h-blob with length ≥ *L*_min_, defined as *H*_max_. SNPs where neither allele reside in an h-blob with this minimum length are discarded from further analysis. The expected heterozygosity of a SNP with frequency *v* (for either the reference or alternative allele) is defined as Θ = 2*v*(1 − *v*). Define the blob length at *H*_max_ as *L*^⋆^. We bin SNPs into four *H*_max_ bins of width Δ*H*_max_ = 0.25 and compute the mean SNP expected heterozygosity within each bin. The null distribution for each bin is computed based on *R* = 5, 000 random permutations of the SNP heterozygosity among the input SNP set. The preceding procedure is tabulated for nSNPs and dSNPs separately.

### Identification of aggregating proteins

There are 28 proteins within the dataset that are annotated as involved in formation of extracellular amyloid deposits or intracellular inclusions with amyloid-like characteristics: P02647, P06727, *P02655, P05067, Q99700, P61769, P01258, *P17927, P07320, P01034, *P35637, P06396, Q9NX55, P10997, P08069, *P01308, P02788, *P61626, P10636, Q08431, P01160, P04156, P11686, P37840, P00441, *Q13148, *Q15582, P02766. For analyses involving aggregation, the uniprot dataset was divided into the SNPs within these 28 proteins (“Aggregating Proteins”) and all other proteins (“non-Aggregating Proteins”).

### Gene pathway and ontology enrichment tests

The gene-ontology enrichment is performed using the g:Profiler web service ***Raudvere et al. (2019***). We use the g:GOSt functional annotation enrichment tool. Statistical p-values are adjusted for multiple-testing and ontology overlap using the g:Profiler algorithm “g:SCS” on a user significance threshold of 0.05. We use custom backgrounds over annotated genes only. The background used for nSNPs is all proteins containing an nSNP and the background for dSNPs is all proteins containing a dSNP. The ontology databases tested for enrichment are the g:Profiler databases as of 2019: GO Molecular Function (GO MF), GO Biological Process (GO BP), GO Cellular Component (GO CC), Kyoto Encylopedia of Genes and Genomes pathways (KEGG), Reactome pathways (REAC), WikiPathways (WP), TRANSFAC (TF), miRTarBase (MIRNA), the Human Protein Atlas (HPA), the Comprehensive Resource of Mammalian protein complexes (CORUM), and the human phenotype ontology (HP).

### Statistical Information

Throughout this paper, enrichment of dSNPs relative to nSNPs is tested using the Binomial distribution for single category tests, and the multinomial distribution for multi-category tests. In general, the number of tests is the number of dSNPs, with the per-test probability of “success” given by the nSNP rate. The probability of having a number of successes at least as extreme as the observed number of dSNPs is the reported (two-sided) p-value. See the above subsections for detailed numbers for each of the enrichment tests.

For the population frequency signature of differential selection, we use permutation tests to quantify the probability of having observed a mean heterozygosity per SNP hydrophobicity bin, described in detail above. The gene ontology and pathway enrichment tests use the g:SCS algorithm for correcting p-values for multiple-tests in the context of overlapping hierarchy of non-independent tests.

### Computational packages used

All computations were done in Python 3.6 using the *numpy, scipy*, and *pandas* packages. All figures are made using the Python *matplotlib* package. The blobulation software is available at https://github.com/BranniganLab/blobulator.

## Acknowledgments

The authors acknowledge the Office of Advanced Research Computing (OARC) at Rutgers, The State University of New Jersey for providing access to the Amarel cluster and associated research computing resources that have contributed to the results reported here. The authors are also grateful to Ms. Kaitlin Bassi and Mr. Connor Pitman for useful discussions and comments on the manuscript, and to Dr. Cameron Abrams for coining the term “blobulation” to describe our algorithm.

## Author Contributions

R.L. and G.B. established the blobulation approach. R.L. wrote the blobulation software and performed enrichment analyses. M.H. ran population-level analyses. All authors contributed to design of the study, discussion of results, generation of figures, and writing of the manuscript.

