## Supplemental Text and Figure for "Contiguously-hydrophobic sequences are functionally significant throughout the human exome"

**Dataset 1:** Enrichment of dSNPs relative to nSNPs in h blobs for various hydrophobicity cutoffs and blob lengths. These data correspond to Fig 2b of the Main Text (blobulation panel).

**Dataset 2: GO enrichment for nSNPs in highly hydrophobic h-blobs** Gene set pathway and ontology enrichment test results using g:Profiler (Raudvere *et al*, 2019) for target proteins containing nSNPs with maximum hydrophobicity $H_{\text{max}} \geq 0.75$ ($n = 635$), compared to a background of all proteins containing an nSNP ($n = 10,406$). See Methods for details.

**Dataset 3: GO enrichment for dSNPs in highly hydrophobic h-blobs** Similar to Table S2 but for dSNPs, where there are $n = 179$ target proteins with $H_{\text{max}} \geq 0.75$ compared against a background of $n = 1,803$ proteins that contain a dSNP.

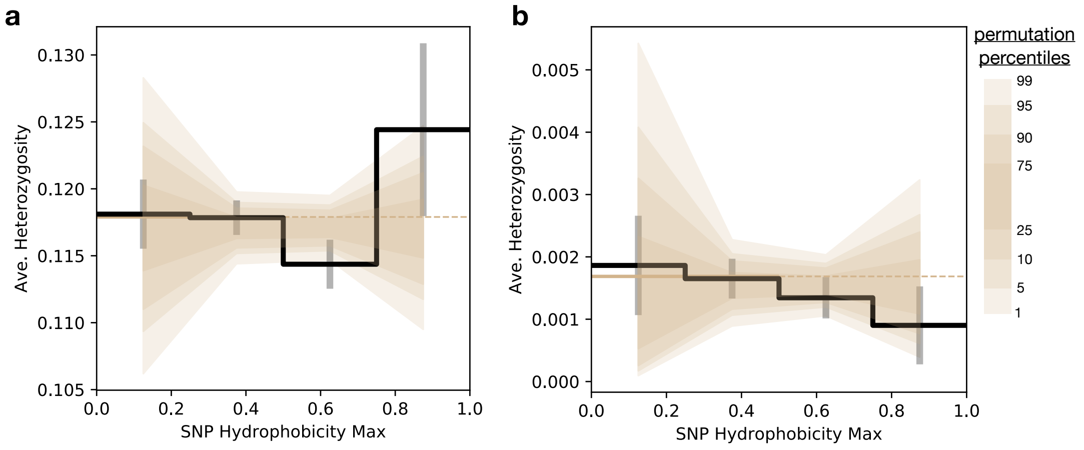


**Fig S1. Heterozygosity of SNPs in East Asians as a function of blob hydrophobicity**. Similar to Figure 2 but for the gnomAD East Asian population cohort. See Methods for details.
